# OSCAR: an open-source controller for animal research

**DOI:** 10.1101/2023.02.03.527033

**Authors:** Evan M. Dastin-van Rijn, Elizabeth Sachse, Francesca Iacobucci, Megan Mensinger, Alik S. Widge

## Abstract

Operant animal behavior training and monitoring is fundamental to scientific inquiry across fields necessitating evaluation via controlled laboratory tasks. However, current commercial and open-source systems enforce particular hardware and software, limiting reproducibility and technique and data sharing across sites. To address this issue, we developed OSCAR: an open-source controller for animal research that enables flexible control of a variety of industry standard hardware with platform-independent software. OSCAR offers millisecond latency with a flexible array of inputs and outputs at a fraction of the cost of commercial options. These features position OSCAR as a valuable option for improving consistency of behavioral experiments across studies.

## Introduction

Operant animal behavior training and monitoring is fundamental to scientific inquiry across fields [1]. In many cases, a behavior of relevance, or its neural substrate, is best studied through a controlled laboratory task. These tasks require tight integration of the hardware components with which animals interact and the overarching software that coordinates these components to elicit desired behaviors. There is a plethora of options for systems to facilitate behavioral tasks, from commercial solutions to open-source developers [2–4]. However, hardware and software from each manufacturer are inextricably linked, limiting reproducibility and technique and data sharing across sites. To address this issue, we developed OSCAR: an open-source controller for animal research that enables flexible control of a variety of industry standard hardware with platform-independent software.

## Methods (Design)

### Considerations

- Compatibility with voltage and current constraints of standard commercial hardware
- Minimal cost
- Simple assembly
- Firmware written in an open-source language and framework
- Analog capabilities
- Sufficient digital options for standard behavioral tasks
- Separation between task code and hardware control
- Compatible with arbitrary task software
- Millisecond scale latency

### Hardware Overview

We developed OSCAR to be a bridge between a simple microcontroller and industry standard −28V hardware. OSCAR is based on the Arduino Mega 2560 microcontroller board (www.store.arduino.cc/products/arduino-mega-2560-rev3), which utilizes an 8 bit ATmega 2560 processor running at 16 MHz. The Mega was chosen for the large number of available inputs/outputs but similar microcontrollers could be used instead for future iterations.The foundation of OSCAR is optical isolation of the 5 V microcontroller from a −28 V supply. Each OSCAR supports 32 digital outputs and 8 digital inputs connected to two 2×20 pin ribbon connectors. Each digital output can provide a maximum of 100 mA of current. Larger currents (up to 3 A) can be driven by using the output as a command signal for a relay connected directly to the −28 V power bus on the external component. Each ribbon connector also includes an audio feedthrough from panel mounted AUX connectors. Two additional terminal blocks (2×4 connections) are provided; one for general purpose input/output (GPIO) and 10 bit analog inputs (configurable from 0-5V or 0-2.5V and 1-1000 Hz sampling rate) and the other for 16 bit analog outputs (0-2.5V). Our circuit board design was made using KICAD (www.kicad-pcb.org), an open-source printed circuit board (PCB) design program. Schematic, layout, bill of materials, and build instructions are included in the OSCAR Github repository (www.github.com/tne-lab/OSCAR). An overview of the design is shown in Figure 1.

**Figure 1:**
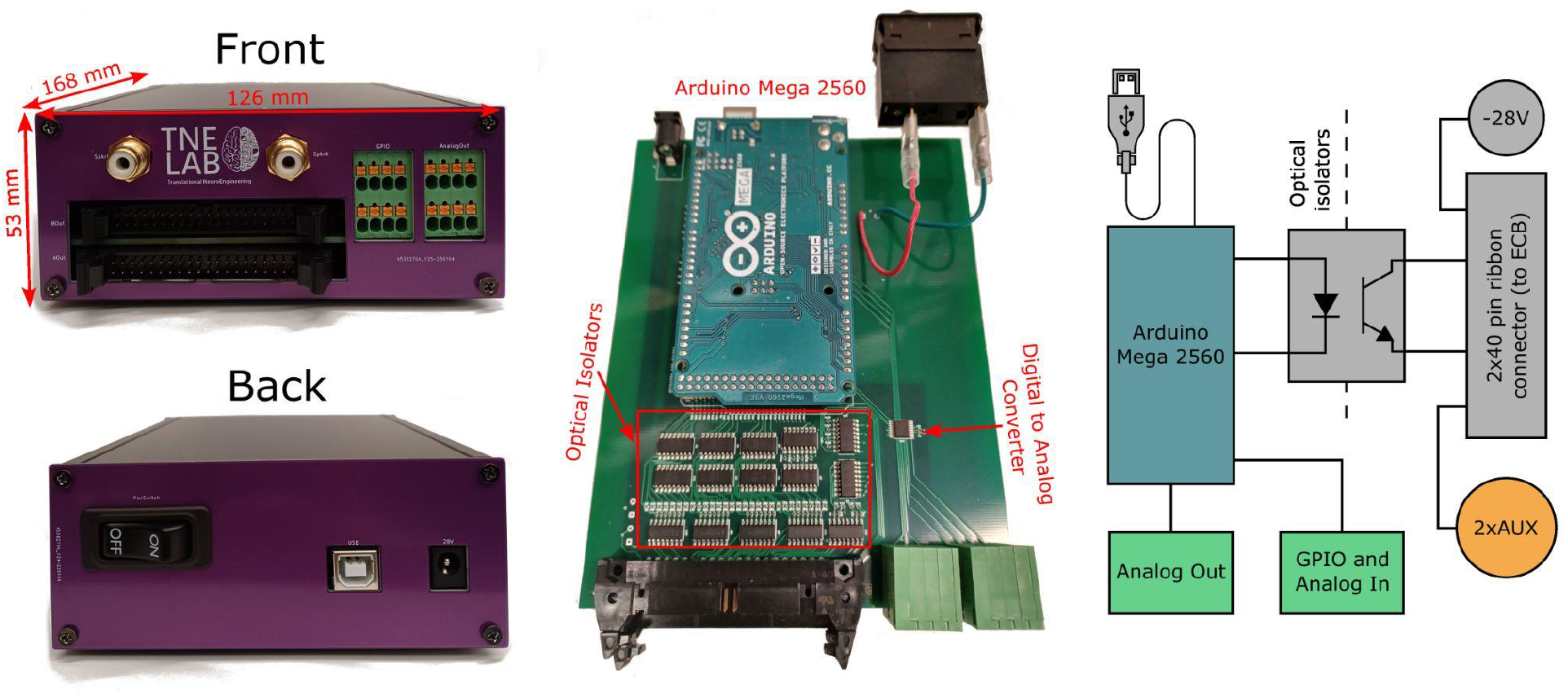
OSCAR layout. OSCAR has a compact footprint, measuring 168×126×53 mm. The front panel (top left) includes two 40-pin ribbon connectors to interface with environment connection boards (ECBs). Two 4×2 terminal blocks are included, one for GPIO connections and the other for analog outputs. Two AUX connectors are additionally provided for audio feedthrough. On the back (lower left) is the USB connector (which also powers the Arduino) and a barrel connector for −28 V input. A circuit breaker is included for the −28 V connection. The internal circuit board (middle) consists of an Arduino Mega2560 which interfaces with optical isolators to switch the −28 V connections. A digital-to-analog converter (DAC) coordinates analog outputs. An overview of the system is shown on the right.

### Software

OSCAR’s software is written in C++ using the Arduino development environment. We provide Arduino-compatible firmware for interfacing with OSCAR’s various inputs and outputs over a serial connection. Serial commands are composed of formatted bit sequences to reduce overhead from information transfer between OSCAR and the host computer. Each command (CB) begins with a sequence of three bits specifying the command type followed by any necessary bits for component addresses, output values, or type. To be compatible with a serial interface, each command is packed into a whole number of bytes with any extra bits being ignored. Further details for the existing commands are included in Table 1. The decision was made to separate hardware control on the microcontroller from task logic to enable compatibility with arbitrary task software and computationally intensive functionality that would be infeasible on the microcontroller. A Python server interface is provided which handles the serial processing, should it be preferable for a user to handle OSCAR communication in a separate process with a local socket connection facilitated by a Router-Dealer ZeroMQ protocol. The server can communicate with multiple OSCARs simultaneously. Instead of binary, server commands begin with the name of the command followed by the OSCAR index the command is related to then any numeric quantities related to the command. All commands to and from the server are separated with line breaks (“\n”).

**Table 1:**
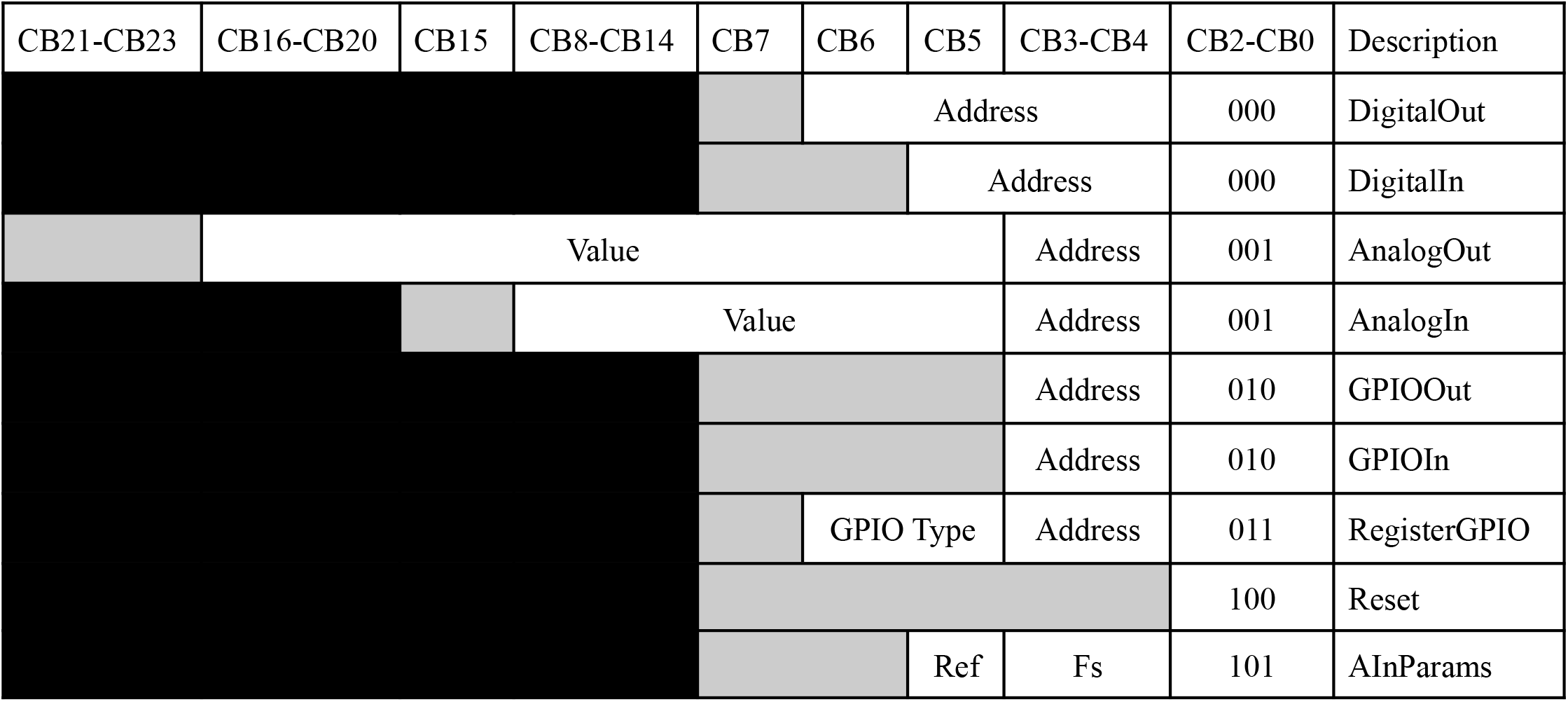
Bit sequences for available OSCAR serial commands. Each command has an ID (separate for input and output commands) and additional bits for relevant instructions. All commands are little-endian.

### Environment Connection Boards

To support interfacing with a variety of components from different manufacturers, users can connect OSCAR to custom environment connection boards (ECBs). Each ECB can connect to one or both of the 40 pin ribbon connectors to break out into any custom connectors that might be required for particular behavioral hardware. Schematics for an example ECB with RJ12 breakouts for compatibility with Coulbourn Instruments hardware is provided (Figure 2).

**Figure 2:**
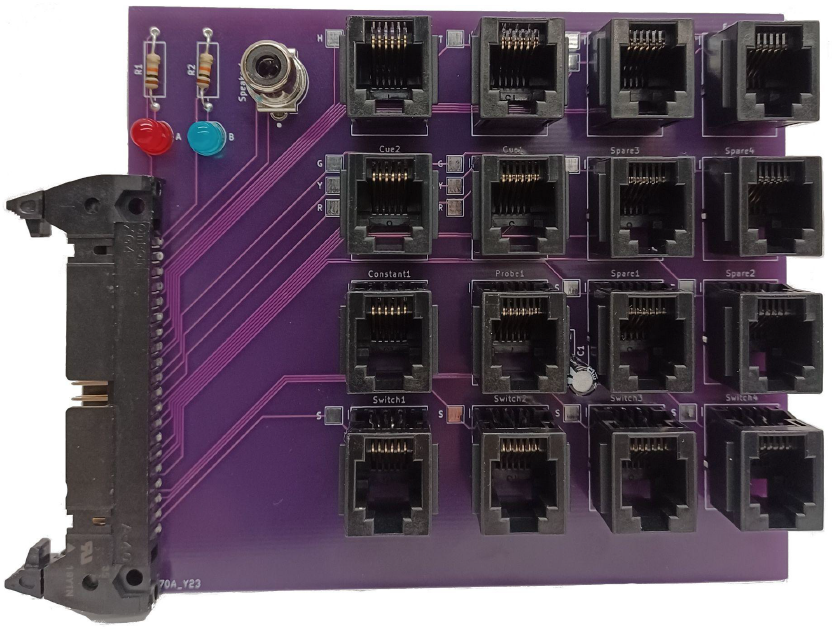
Example Environment Connection Board. This figure shows an example of a potential environment connection board (ECB) that would be compatible with legacy hardware from Coulbourn Instruments. Each input/output from the 40 pin ribbon connector is broken out into RJ12 connectors for use with Coulbourn components.

### Comparison to competitors

OSCAR’s design compares favorably against alternatives (Table 2). It is similarly priced to open-source systems like Pycontrol [2] but an order of magnitude less expensive than the commercial hardware it is designed to replicate and interact with. However, the existing firmware for OSCAR is optimized for relaying serial commands to and from the device to coordinate with external task code rather than directly running tasks on the Arduino. This leads to slightly higher latencies compared to an embedded system, like Pycontrol. Custom firmware could be written for this purpose should a user require OSCAR’s capabilities with more refined latencies.

**Table 2:** Comparison of OSCAR vs. other related hardware. Cost was obtained as the complete cost of the bill of materials for OSCAR or the quoted price from the distributor for other hardware. Extension modules were not included in this assessment. Current in parentheses is the maximum total output from the corresponding device (which can be used to drive a single component dependending on the configuration). * Pycontrol connections are grouped into 6 RJ45 and 4 BNC connectors. ^†^Panlab connections are grouped into 8 RJ12 connectors.

|  | OSCAR | Pycontrol | BPod r2+ | Panlab | Med Associates | Lafayette Instruments |
| --- | --- | --- | --- | --- | --- | --- |
| Digital Outputs | 32 (+4) | 18* (shared) | 5 (+4 shared) | 16<sup>†</sup> (shared) | 16 | 30 |
| Digital Inputs | 8 (+4) | 18* (shared) | 4 (shared) | 16<sup>†</sup> (shared) | 8 | 16 |
| Analog Inputs | 4 | 6* (shared) | 4 (shared) | 0 | 0 | 0 |
| Analog Outputs | 4 | 4* (shared) | 4 (shared) | 0 | 0 | 0 |
| Component operating voltage | -28 V | 12 V | 12 V | -28 V | -28 V | -28 V |
| Max current draw | 100 mA (3 A) | 200 mA (600 mA) | 250 mA | Unavailable | (3 A) | (1.25 A) |
| Cost (USD) | \$162 | \$256 | \$1745 | Unavailable | ~\$2600 | ~\$1600 |
| Task on controller | N (with standard firmware) | Y | Y | N | N | N |
| Open-source | Y | Y | Y | N | N | N |

## Results

### Benchmarking

To benchmark OSCAR, we ran two separate, 30 s latency tests for the digital and analog components. This latency reflects the time to detect the input. For the digital test, we provided a 50 Hz square pulse train to a digital input and replicated the pulse train with a digital output. For the analog test, we provided a 50 Hz, 2.5 V sine wave to the analog input and replicated the wave with an analog output. Both test sequences were provided and recorded by a National Instruments USB-6343 DAQ (250 kHz sampling rate) using scripts written in MATLAB 2021b. Digital tests were performed under a ‘low load’ condition where the board was processing no other connections and a ‘high load’ condition where two 1000 Hz analog inputs were simultaneously monitored. In the ‘low load’ condition, response latency was 1453 ± 224 μs (mean ± SD) (Figure 3A) while in the ‘high load’ condition it was 1559 ± 319 μs (mean ± SD) (Figure 3B). The longest latency recorded was 2.532 ms. For the analog test, response latency was 1190 ± 334 μs (mean ± SD) (Figure 3 C-D). Errors in the analog test are dependent on the resolution of the bidirectional digital/analog conversion and frequency of the signal of interest.

**Figure 3.**
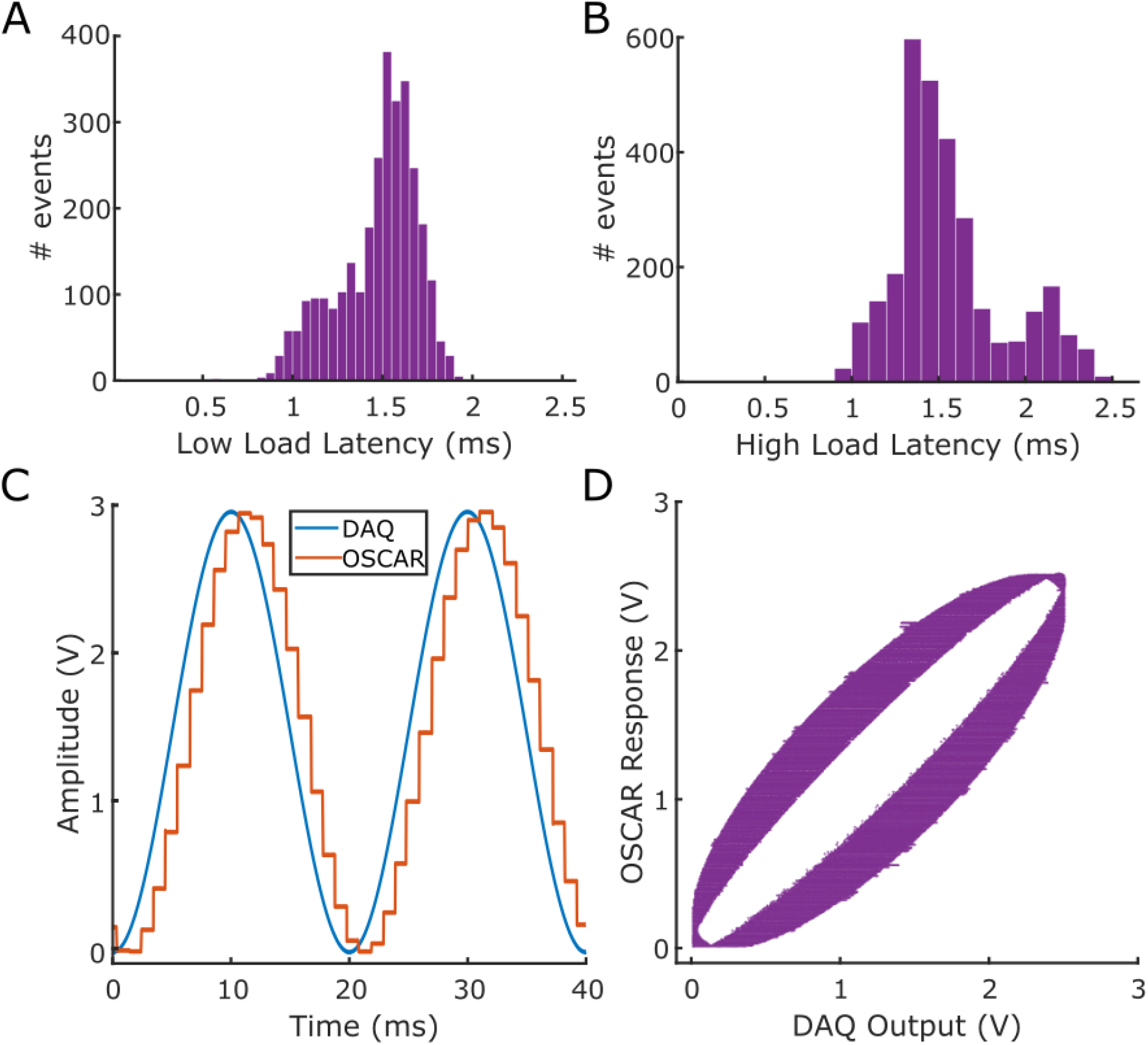
Latency benchmarking. **(A)** Distribution of latencies for OSCAR to respond to a change in a digital input by changing the level of a digital output. **(B)** Same as **(A)** but under a high load condition. **(C)** Comparison of DAQ signal (blue) to OSCAR response after replicating an analog input with an analog output. **(D)** Scatter of OSCAR input amplitude against DAQ output amplitude.

### Sample Use Case

We include an example of how OSCAR has been used in practice with a standard task in an operant chamber: the 5-choice serial reaction time task (5-CSRTT) [5]. In this task, a rat must detect a brief flash of light presented pseudorandomly in one of five nose-poke ports and respond to the stimulus location by poking the corresponding port to trigger a reward delivered to a receptacle on the opposing wall (Figure 4A). We used a standard operant box developed by Coulbourn Instruments with 5 IR nose pokes with white LEDs, a food receptacle with an LED, a motorized reward pellet delivery system, and house lights. All of these components were connected to a single OSCAR via two of the previously described ECBs via RJ12 connectors. The task code itself was implemented using an in development Python framework (www.github.com/tne-lab/py-behav-box-v2). The example session shown demonstrates that a subject could elicit the full scope of available behaviors (Figure 4B) and had typical reaction time distributions (Figure 4C) [5].

**Figure 4.**
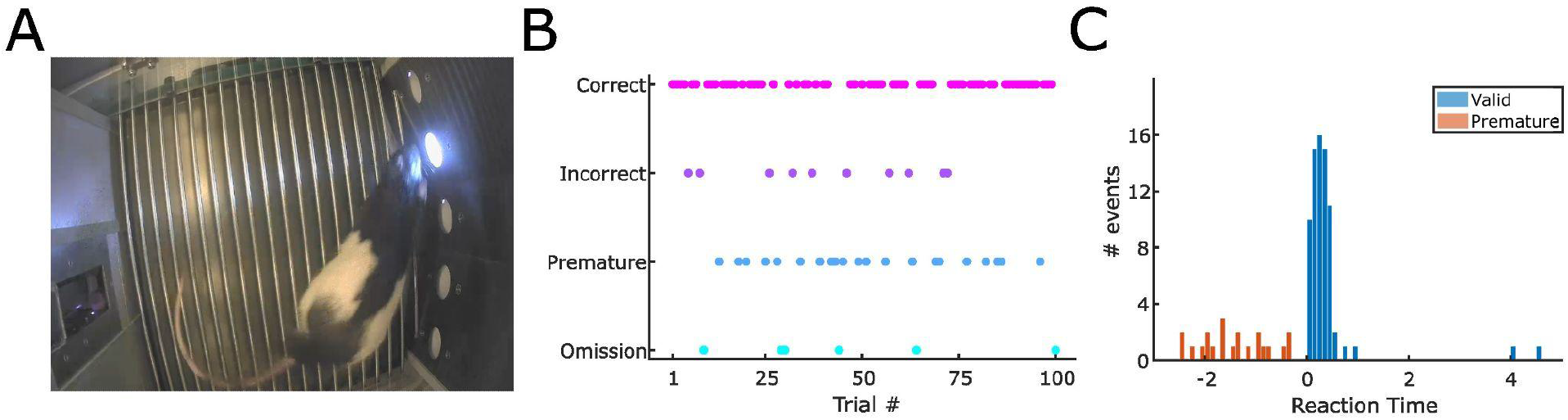
OSCAR facilitates a five choice serial reaction time task. **(A)** Example image of a rat performing the task in a chamber coordinated by OSCAR. **(B)** The full scope of possible behaviors (omitted, premature, incorrect, and correct responses) was observed throughout the task. **(C)** Distributions of reaction times for premature and valid responses.

## Conclusions

We have introduced OSCAR, an inexpensive yet flexible open-source controller for behavioral neuroscience experiments. OSCAR is designed to be compatible with commercial components at a fraction of the cost of their commercial, proprietary controllers. It offers millisecond latency with a flexible array of inputs and outputs. OSCAR’s low cost and open-source framework enables researchers to easily share and execute behavioral tasks across labs regardless of the specific hardware they might use.

## Author Contributions

ED and AW conceived of the need for the system. ED designed OSCAR with assistance from ES and AW. FI and ED designed the environmental connection boards. MM ran the five choice serial reaction time task. ED made the figures and wrote the manuscript with input from ES, FI, MM, and AW.

## Bibliography

[1] J.W. Krakauer, A.A. Ghazanfar, A. Gomez-Marin, M.A. MacIver, D. Poeppel, Neuroscience Needs Behavior: Correcting a Reductionist Bias, Neuron. 93 (2017) 480–490. https://doi.org/10.1016/j.neuron.2016.12.041.

[2] T. Akam, A. Lustig, J.M. Rowland, S.K. Kapanaiah, J. Esteve-Agraz, M. Panniello, C. Márquez, M.M. Kohl, D. Kätzel, R.M. Costa, M.E. Walton, Open-source, Python-based, hardware and software for controlling behavioural neuroscience experiments, ELife. 11 (2022) e67846. https://doi.org/10.7554/eLife.67846.

[3] N. Buscher, A. Ojeda, M. Francoeur, S. Hulyalkar, C. Claros, T. Tang, A. Terry, A. Gupta, L. Fakhraei, D.S. Ramanathan, Open-source raspberry Pi-based operant box for translational behavioral testing in rodents, Journal of Neuroscience Methods. 342 (2020) 108761. https://doi.org/10.1016/j.jneumeth.2020.108761.

[4] K. Gurley, Two open source designs for a low-cost operant chamber using Raspberry Pi™, Journal of the Experimental Analysis of Behavior. 111 (2019) 508–518. https://doi.org/10.1002/jeab.520.

[5] A. Bari, J.W. Dalley, T.W. Robbins, The application of the 5-choice serial reaction time task for the assessment of visual attentional processes and impulse control in rats, Nat Protoc. 3 (2008) 759–767. https://doi.org/10.1038/nprot.2008.41.

